# BZLF1 interacts with the chromatin remodeler INO80 promoting escape from latent infections with Epstein-Barr virus

**DOI:** 10.1101/317354

**Authors:** Marisa Schaeffner, Paulina Mrozek-Gorska, Anne Woellmer, Takanobu Tagawa, Alexander Buschle, Nils Krietenstein, Corinna Lieleg, Philipp Korber, Wolfgang Hammerschmidt

## Abstract

A hallmark of Epstein-Barr virus (EBV) infections is its latent phase, when all viral lytic genes are repressed. Repression results from a high nucleosome occupancy and epigenetic silencing by cellular factors such as the Polycomb repressive complex 2 (PRC2) and DNA methyltransferases that respectively introduce repressive histone marks and DNA methylation. The viral transcription factor BZLF1 acts as molecular switch to induce the transition from the latent to the lytic or productive phase of EBV’s life cycle. It is unknown how BZLF1 can bind to the epigenetically silenced viral DNA and whether it directly reactivates the viral genome through chromatin remodeling. We addressed these fundamental questions and found that BZLF1 binds to nucleosomal DNA motifs both *in vivo* and *in vitro*, a property characteristic of *bona fide* pioneer factors. BZLF1 co-precipitates with cellular chromatin remodeler ATPases, and the knock-down of one of them, INO80, impaired lytic reactivation and virus synthesis. We conclude that BZLF1 reactivates the EBV genome by directly binding to silenced chromatin and recruiting cellular chromatin remodeling enzymes, which implement a permissive state for viral transcription. BZLF1 shares this mode of action with a limited number of cellular pioneer factors, which are instrumental in transcriptional activation, differentiation, and reprogramming in all eukaryotic cells.

## INTRODUCTION

Eukaryotic DNA binding sites are often not accessible to their cognate factors because the sites lie within epigenetically silent chromatin and are occupied by nucleosomes. Nucleosomes at binding sites constitute a physical barrier to transcription factors because their binding is often structurally incompatible with DNA wrapped around the histone octamer. Access to nucleosomal sites may be achieved through cooperative and simultaneous binding of several transcription factors that outcompete the histone octamer (Adams and Workman, 1995; Mirny, 2010). Alternatively, one class of transcription factors, termed pioneer factors (Magnani et al., 2011b; Zaret and Carroll, 2011; Cirillo et al., 1998; Cirillo et al., 2002), can bind their target sequences even on nucleosomal DNA and in silent chromatin and establish competence for gene expression through chromatin remodeling (Zaret and Mango, 2016 for a recent review). Pioneer factors either open chromatin directly through their binding or recruit chromatin modifiers and ATP-dependent chromatin remodeling enzymes that open chromatin to allow access for the transcription machinery (Bartholomew, 2014; Clapier and Cairns, 2009; Längst and Manelyte, 2015). Such pioneer factors play key roles in hormone-dependent cancers (Jozwik and Carroll, 2012), embryonic stem cells and cell fate specification (Smale, 2010; Drouin, 2014), and cellular reprogramming (Iwafuchi-Doi and Zaret, 2014; Soufi et al., 2015). Currently, two- to three-thousand sequence-specific DNA-binding transcription factors in human cells are known (Lander et al., 2001; Venter et al., 2001), but only about a dozen are characterized as pioneer factors.

Certain pioneer factors have peculiar structural characteristics that explain binding to nucleosomal DNA. For example, the winged-helix DNA binding domain of the paradigm pioneer factor FoxA structurally resembles the linker Histone H1, disrupts inter-nucleosomal interactions, opens chromatin, and enhances *albumin* expression in liver cells (Cirillo et al., 2002; Sekiya et al., 2009). How many other pioneer factors bind to nucleosomal DNA is less well understood, but some directly target partial DNA motifs displayed on the nucleosomal surface (Soufi et al., 2015). Subsequently, most pioneer factors recruit chromatin remodelers to their binding sites, which open silent chromatin and regulate cell-type specific gene expression (Magnani et al., 2011a; Mayran et al., 2015).

In eukaryotic nuclei, chromatin remodelers mediate the dynamics of nucleosome arrangements and participate in most DNA-dependent processes (Längst and Manelyte, 2015 for a recent overview). They bind to nucleosomes and convert the energy of ATP hydrolysis into the movement, restructuring, or ejection of histone octamers depending on the remodeler. Remodelers are categorized according to their ATPase subunit into four major (SWI/SNF, ISWI, INO80, and CHD) and several minor families and further differentiated by their associated subunits. This range of features reflects specialized functions found in their domains/subunits that mediate direct interactions with modified histones, histone variants, DNA structures/sequences, RNA molecules, and transcription factors. The human genome encodes 53 different remodeler ATPases (Längst and Manelyte, 2015), which are highly abundant chromatin factors with roughly one remodeling complex per ten nucleosomes (Längst and Manelyte, 2015).

The Epstein-Barr Virus (EBV) infects more than 95 % of the adult population worldwide with a lifelong persistence in human B cells. The key to EBV’s success lies in its ingenious multipartite life, which relies on different epigenetic states of viral DNA (Woellmer and Hammerschmidt, 2013). Initially, EBV establishes a latent infection in all cells it infects (Hammerschmidt, 2015; Kalla et al., 2012). Viral latency is characterized by an epigenetically silenced EBV genome that prevents expression of all lytic viral genes, but usually spares a small set of so-called latent viral genes that remain active. Cellular factors, e.g. the Polycomb repressive complex 2 (PRC2) and DNA methyltransferases respectively introduce repressive histone marks and 5-methyl cytosine residues into viral DNA, which ensure the repressed state of all viral lytic genes (Ramasubramanyan et al., 2012; Woellmer et al., 2012).

BZLF1 is the viral factor that acts as a molecular switch, induces the lytic, productive phase of EBV *de novo* synthesis, and hence abrogates transcriptional repression of viral lytic genes (Countryman and Miller, 1985; Countryman et al., 1987; Chevallier-Greco et al., 1986; Takada et al., 1986). BZLF1 binds methylated EBV DNA sequence-specifically (Kalla et al., 2012; Bergbauer et al., 2010; Bhende et al., 2004), but if and how it overcomes epigenetically repressed chromatin is not known.

BZLF1 binds to two classes of BZLF1 responsive elements (ZREs): one class contains a DNA sequence motif reminiscent of the canonical AP-1 binding site, the other class contains a sequence motif with a CpG dinucleotide, which must carry 5-methyl cytosine residues for efficient BZLF1 binding (Bergbauer et al., 2010; Bhende et al., 2004; Flower et al., 2011; Karlsson et al., 2008). Binding of BZLF1 to viral chromatin induces the loss of nucleosomes at certain but not all ZREs with higher than average nucleosome densities (see Figs. 2 and 3 in ref. Woellmer et al., 2012). This study did not determine whether the initial binding of BZLF1 and loss of nucleosomes are simultaneous events or occur sequentially nor did it identify the molecular mechanisms that underlie these events.

**Fig. 1.**
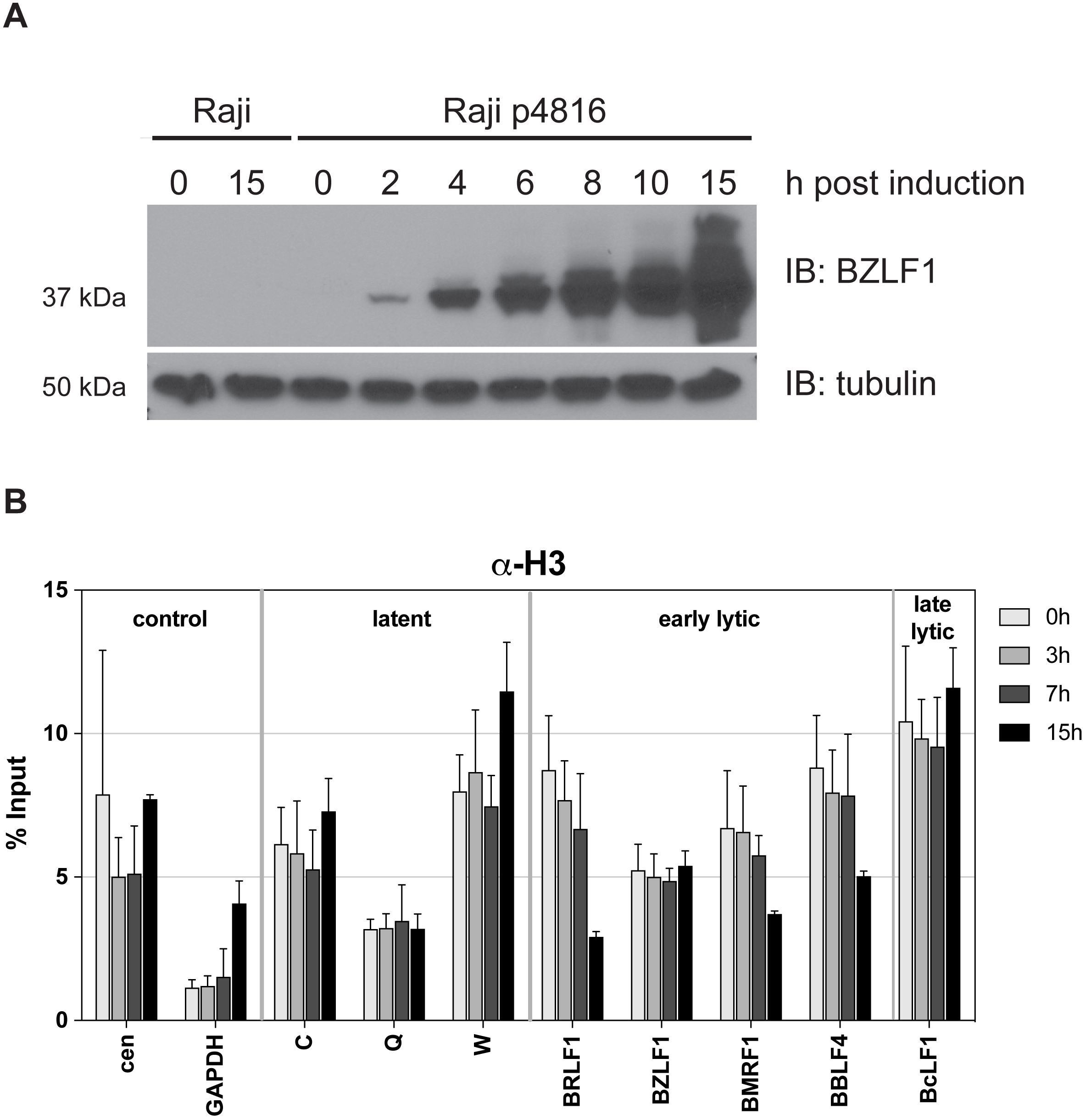
BZLF1 expression precedes H3 loss at promoter sites of early lytic genes in lytically induced Raji cells. **(A)** The kinetics of BZLF1 protein expression in parental and Raji p4816 cells was analyzed by immuno blotting (IB) with a BZLF1-specific antibody (top panel) at the indicated time points (hours post induction). Immunodetection of tubulin (bottom panel) served as loading control. **(B)** Chromatin immunoprecipitation directed against histone H3 (Abcam, #1791) of lytically induced Raji p4816 cells at the indicated time points of induction. Mean and SD from three independent experiments are shown. Primer information can be found in Tab. S2.

**Fig. 2.**
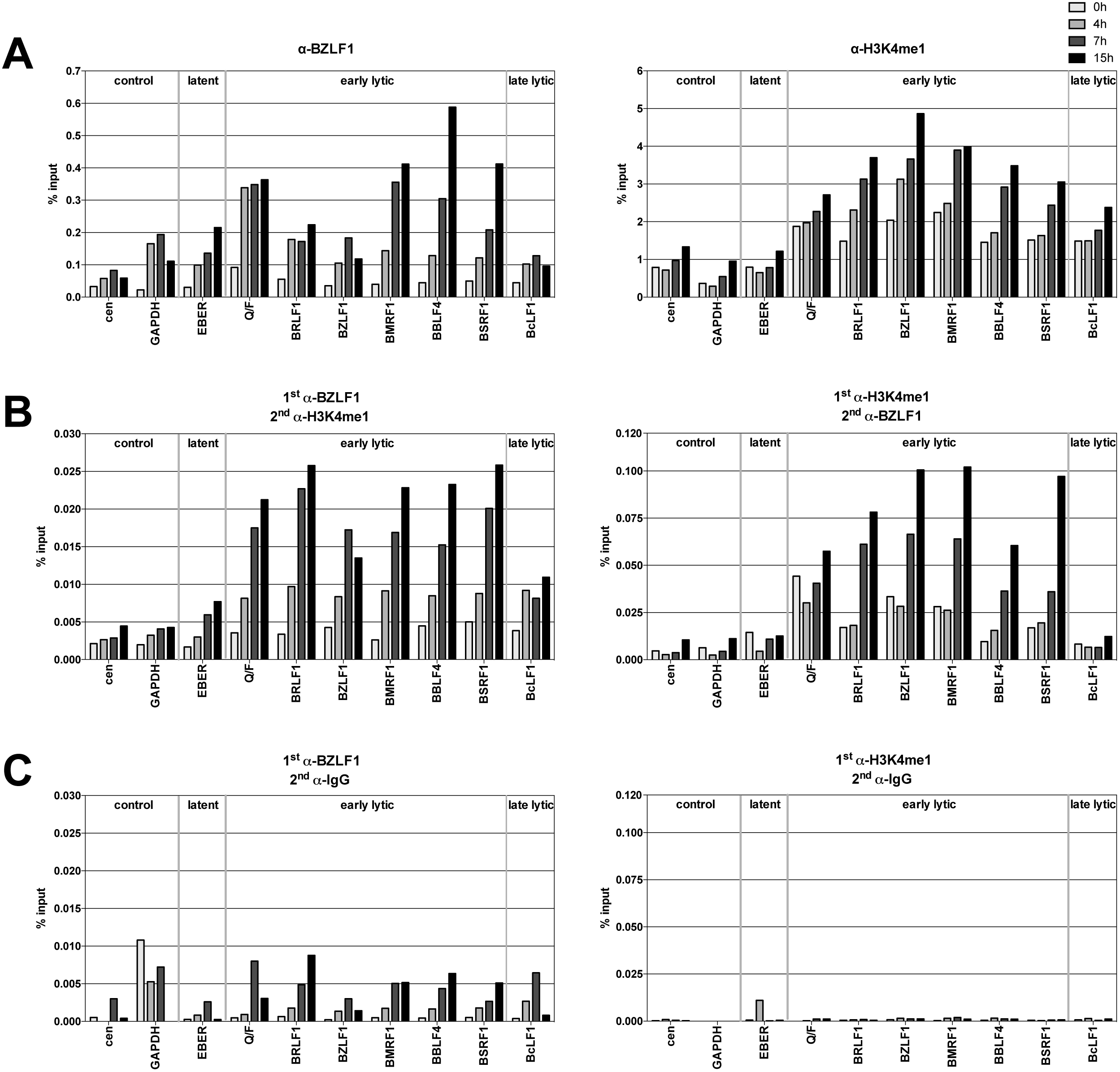
BZLF1 and histone octamers co-occupy promoter sites of early lytic genes *in vivo*. **(A)** qPCR data of ChIP experiments with Raji p4816 cells are shown. Antibodies directed against BZLF1 or H3K4me1 and primer pairs specific for the indicated human (cen, GAPDH) or viral loci were used. Chromatin in the ChIP experiments had been sheared to a size of approximately 150 bp corresponding to mononucleosomal-sized DNA fragments. Primer information can be found in Tab. S2. Mean values of three independent experiments are provided. **(B)** As panel A, but ReChIP experiments with sequential use of two different antibodies against either BZLF1 or H3K4me1. Right and left panel differ in the order of the antibodies used (indicated on top of the panels). Mean values of qPCR analysis of three independent ChIP replicates are provided. **(C)** As panel B, but with non-specific IgG antibody as secondary antibody (indicated on top of the panels).

**Fig. 3.**
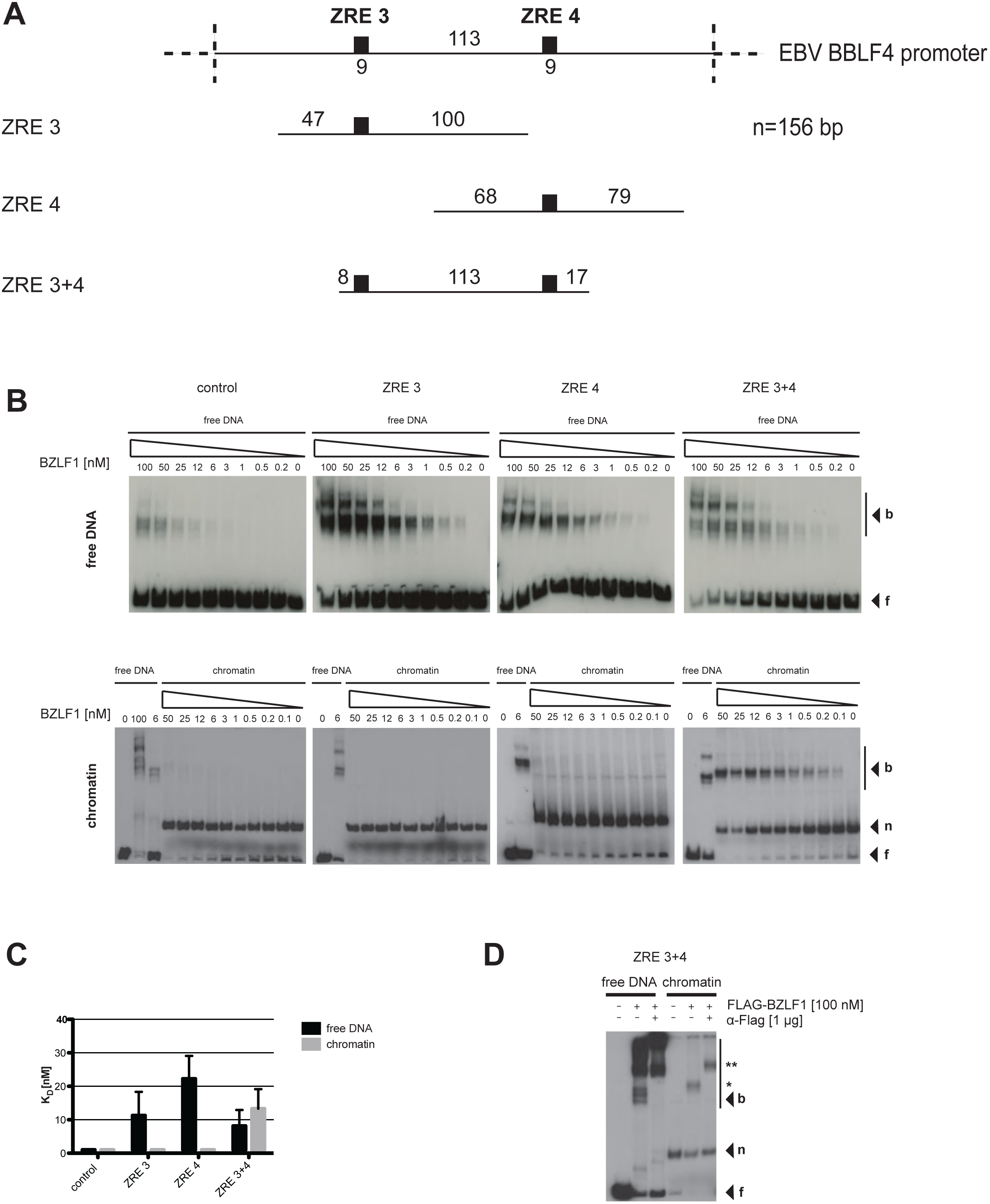
BZLF1 binds mononucleosomes *in vitro*. **(A)** Top: schematics of the relative position of the two BZLF1 responsive elements ZRE 3 and ZRE 4 (black boxes) at the *BBLF*4 promoter. Numbers indicated DNA lengths in bp. Below: schematics of three different 156 bp fragments encompassing the indicated ZREs. Both ZREs contain a CpG motif that must be methylated for BZLF1 binding. **(B)** Electromobility shift assays (EMSAs) for binding of BZLF1 (at indicated concentration) to DNA fragments as in panel A or to an EBV control region without ZRE that is not bound by BZLF1. DNA was either free or assembled into mononucleosomes by salt gradient dialysis (chromatin). The migration positions of free DNA (f), mononucleosomes (n), or the complexes with BZLF1 (b) are indicated on the right of the autoradiographs. **(C)** Quantification of equilibrium dissociation constants (K_D_) from EMSA experiments as in panel B. If error bars are provided, the average and standard deviation of three independent experiments are shown. **(D)** EMSA (“super shift assay”) with FLAG-tagged BZLF1 (FLAG-BZLF1), free or mononucleosomal ZRE 3+4 DNA and anti-FLAG (α-FLAG) antibody as indicated. The migration positions of the FLAG-BZLF1/mononucleosome and the FLAG-BZLF1/α-FLAG /mononucleosome complexes are indicated on the right by one (^*^) and two asterisks (^**^), respectively.

Here we report that the viral factor BZLF1 acts like a pioneer transcription factor. BZLF1 binds mononucleosomal DNA in repressed lytic promoters *in vivo* and binds to nucleosome core particle DNA *in vitro*. BZLF1 interacts with two different chromatin remodelers and likely recruits them to epigenetically repressed viral chromatin. Co-precipitations identify the transcriptional activation domain of BZLF1 as interacting with the remodeler ATPase INO80. A knock-down of INO80, but not of the remodeler ATPase SNF2h, reduces activation of early lytic viral genes and virus *de novo* synthesis, suggesting that INO80-containing remodeler complexes are critical for viral reactivation.

## MATERIALS AND METHODS

### Cells

Raji, THP-1, and HEK293 cells (Pulvertaft, 1964; Berges et al., 2005; Graham et al., 1977) were maintained in RPMI 1640 medium (Thermo Fisher Scientific) supplemented with 10 % fetal calf serum (FCS, Bio&Sell), 1 % penicillin-streptomycin (Thermo Fisher Scientific), 1 % sodium pyruvate (Thermo Fisher Scientific), and 100 nM sodium selenite (Merck) at 37 °C and 5 % CO_2_. 293T cells were kept in DMEM (Thermo Fisher Scientific) including the supplements mentioned above. Raji p4816, p5693 and p5694 cells were kept under constant selection with 1 μg/ml puromycin. The HEK293-based cell line for the production of recombinant wild-type 2089 EBV stocks was cultivated in fully supplemented RPMI 1640 medium with 100 μg/ml hygromycin (Delecluse et al., 1998).

### Plasmids

Plasmids p4816, p5693 and p5694 (Fig. S1) were constructed as described in Woellmer et al (Woellmer et al., 2012). cDNAs coding for *INO80* and *SNF2h* were cloned in frame with the e*GFP* gene and expressed from the CMV promoter in pEGFP-C1 and pEGFP-N3 (Clontech), respectively. All BZLF1 expression plasmids (aa1-245, aa1-236, aa149-245, and aa175-236) were expressed with a FLAG- and tandem Strep-tag (Gloeckner et al., 2007). The BZLF1 and gp110/BALF4 expression plasmids p509 and p2670 have been described (Hammerschmidt and Sugden, 1988; Neuhierl et al., 2002). The plasmid p6605 expresses a W653Q point mutant of human INO80 with a triple HA tag at its carboxy-terminus under the control of the CMV promoter.

### Stable transfection and establishment of Raji cells

5×10^6^ Raji cells were suspended in 250 μl Opti-MEM I medium (Thermo Fisher Scientific), 5-10 μg plasmid DNA was added and the cells were incubated on ice for 15 min. Electroporation (gene pulser II instrument, Bio-Rad) was done in 4 mm cuvettes at 230 V and 975 μF. The cells were resuspended with 400 μl FCS, transferred to 5 ml fully supplemented medium as described above, and cultivated at 37 °C for two days. For the establishment of single cell clones, cells were diluted in 96 well cluster plates and cultivated under selection for four weeks. Medium was changed when necessary, and outgrowing cells were expanded. GFP expression was monitored by flow cytometry with a FACSCanto instrument by Becton Dickinson.

### Transient transfection of cell lines

Transfection of DNA into HEK293 cells using polyethylenimine (Polysciences, #24765) was as described (Reed et al., 2006). For protein extracts, 2×10^7^ cells per 13 cm cell culture dish were seeded the day before transfection. Each plate was transfected with 30 μg plasmid DNA.

### Chromatin immunoprecipitation (ChIP) and sequential chromatin immunoprecipitation (ReChIP)

All ChIP experiments were performed in triplicates as described previously (Woellmer et al., 2012) using anti-H3 (Abcam, #1791), anti-H3K4me1 (Abcam, #8895), anti-BZLF1 (Santa Cruz, #17503) antibodies, or control IgG antibody (Millipore, #PP64B). All buffers were supplemented with the cOmplete protease inhibitor cocktail (Roche) and all steps were performed at 4°C if not noted otherwise. Details of the ChIP and ReChIP protocols can be found in the Supplementary Materials and Methods.

Immunoprecipitated DNA was purified with the NucleoSpin Extract II Kit (Macherey-Nagel) according to the manufacturer’s protocol and eluted in 60 μl elution buffer. Samples were analyzed by qPCR with a LightCycler 480 (Roche) instrument.

### Expression of HA-tagged INO80 (W653Q) and ChIP of 2089 EBV HEK293 cells

HEK293 cells were transiently transfected with an expression plasmid encoding an HA-tagged point mutation of INO80 (W653Q). ChIPs were performed as detailed in the Supplementary Materials and Methods.

### Quantitative real time PCR (qPCR)

Immunoprecipitated DNA was quantified with a Roche LightCycler 480 instrument. PCR mixes consisted of template DNA (1 μl), primers (5 pm each), 2x SYBR Green I Master mix (5 μl) in a final volume of 10 μl. The PCR program for qPCR are listed in Tab. S1 in Supplementary Data.

Primer design criteria were as follows: 62 °C annealing temperature, primer efficiency of 2.0 and a single melting peak during Roche LightCycler 480 measurement. Primer synthesis was performed by Metabion (Munich) and sequences are listed in Tab. S2 in Supplementary Data.

Absolute quantifications of the amount of DNA were calculated by comparing the crossing points (Cp) of the unknown sample with a defined standard curve, which encompassed different dilutions of input DNA. The analysis was done automatically with the LightCycler 480 software according to the “second derivative maximum method”. Mean and standard deviation were calculated from three independent biological replicates with one technical replicate each.

### *In vitro* DNA methylation

CpG methylation of plasmid DNA *in vitro* was done with the *de novo* methyltransferase M.SssI (New England Biolabs) and S-adenosyl methionine according to the manufacturer’s recommendations.

### *In vitro* reconstitution of chromatin

*Drosophila* embryo histone octamers were prepared and used for *in vitro* nucleosome reconstitution via salt gradient dialysis according to Krietenstein et al. (Krietenstein et al., 2012).

### Electromobility shift assays (EMSAs)

EMSAs were performed according to Fried and Crothers (Fried and Crothers, 1981). Proteins were purified from HEK293 cells transiently transfected with a FLAG- and tandem Strep-tagged BZLF1 expression plasmid (Bergbauer et al., 2010). Two days post transfection cells from six 13 cm cell culture dishes were pooled and lysed in 10 ml RIPA-buffer (50 mM Tris-HCl pH 8.0, 150 mM NaCl, 1 % Ipegal, 0.5 % DOC, 0.1 % SDS). Lysates were sonicated as above and BZLF1 protein was purified using Strep-Tactin affinity chromatography according to the manufacturer’s recommendations (IBA). All buffers were supplemented with the cOmplete protease inhibitor cocktail (Roche). Free and reconstituted DNA was radioactively labeled with [γ-^32^P] ATP by T4 polynucleotide kinase (NEB) according to manufacturer’s recommendations. For each EMSA reaction, 0.4 nM of radioactively labeled, free or reconstituted DNA was incubated with different concentrations of BZLF1 protein in the presence of 10 mM Tris-HCl pH 7.6, 1 mM MgCl_2_, 60 mM KCl, 3 mg/ml BSA, 1 % glycerol, 1 % Ficoll, 1 mM DTT, 1 μg polydIdC (Roche), and 100 ng calf thymus DNA (Merck) in a total volume of 20 μl for 10 minutes at room temperature. Unbound template was separated from shifted complexes by polyacrylamide gel electrophoresis (5 % (w/v) 29:1 acrylamide/bisacrylamide, 0.5xTBE). Gels were analyzed with the aid of a radioisotope scanner (FLA 5100, Fuji) and quantitation of radioactivity signals was performed with the AIDA program (Raytest). For determination of the equilibrium dissociation constant (K_D_), these data were fitted to the Hill equation with one-site specific binding using the PRISM 6 program (Graphpad).

### Co-immunoprecipitation (Co-IP)

Co-IPs of GFP-fusion or Strep-tag fusion proteins were done with GFP-Trap_A (Chromotek) or Strep-Tactin affinity chromatography using Strep-Tactin sepharose beads (IBA), respectively, according to manufacturer’s recommendations. All buffers contained the cOmplete protease inhibitor cocktail (Roche). Lysis buffers were supplemented with 5 U/μl Benzonase (Merck) and 0.5 μg/μl DNase I (Invitrogen) to exclude nucleic acid mediated interactions.

### Western blotting

Proteins separated by SDS-PAGE were transferred (Mini Trans-Blot Cell, Bio-Rad) onto Hybond ECL membrane (GE Healthcare) at 100 V for 80 minutes and detected with the respective antibodies using the ECL reagent and X ray films (GE Healthcare). Primary antibodies were anti-BZ1 (kindly provided by Elisabeth Kremmer, Helmholtz Zentrum München), anti-SNF2h (Active Motif, #39543), anti-INO80 (Proteintech #18810-1-AP), anti-CHD4 antibody (Abcam #ab70469), anti-GFP (Abcam, #290), and anti-tubulin (Santa Cruz, #23948) and used at appropriate dilutions in blocking buffer (5 g skimmed milk powder in 100 ml PBS-T).

### siRNA knock-down

Transfections were performed with commercial siRNAs directed against transcripts of SNF2h/SMARCA5 (Dharmacon, #E-011478-00) or INO80 (Dharmacon #E-004176-00) or with a random, non-targeting siRNA control pool (Dharmacon #D-001910-10-05) (Fig. S9C). siRNA transfections were done in serum-free Opti-MEM I medium (Thermo Fisher Scientific) in combination with HiPerFect transfection reagent (Qiagen). 2089 EBV HEK293 cells (Delecluse et al., 1998) were transfected with the siRNAs for three days prior to transient transfection of expression plasmids encoding BZLF1 and gp110/BALF4 to induce virus production in the siRNA-treated cells.

### Cell viability assay

siRNA transfected 2089 EBV HEK293 cells were seeded at three different densities into opaque-walled 96-well plate in 100 μl/well. 100 μl of CellTiter-Glo reagent (Promega) was added and mixed for 2 minutes on an orbital shaker to induce cell lysis. The plate was incubated at room temperature for 10 minutes to stabilize the luminescent signal, which was subsequently recorded using a ClarioStar plate reader. Viability of the cells was calculated and expressed as percent of viable cells compared to cells treated with non-targeting siRNA.

### Generation of Raji p4816 cell lines with lentiviral shRNA vectors directed against INO80

Potentially suitable shRNAs were identified using the publicly available web tool siDirect2.0 (http://sidirect2.rnai.jp/). Based on the identified sequences, primers were designed and cloned into the pCDH lentiviral vector (System Biosciences), which was modified and termed p6573 as shown in Figure S9A. The lentiviral vector encodes the red fluorescent protein (DsRed) as a marker gene. Details of the production of lentiviral vectors can be found in the Supplementary Materials and Methods. Raji p4816 cells stably transduced with shRNA_nt (non-targeting) or INO80-specific shRNAs were used in the experiments shown in Figures 7, S9, and S10.

**Fig. 4.**
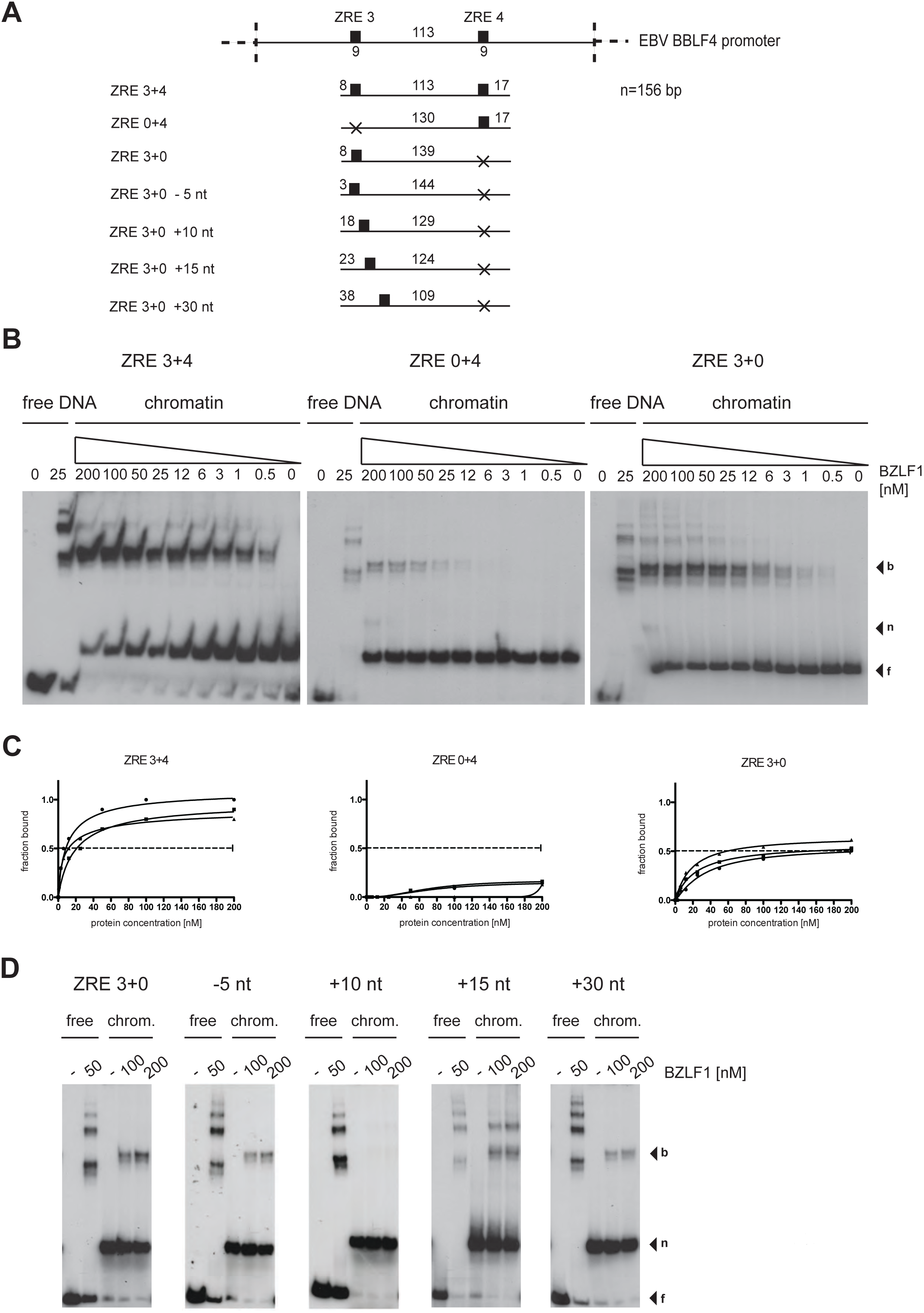
BZLF1 shows cooperative binding to the nucleosomal core *in vitro*. **(A)** Shown are the DNA templates used for the functional analysis of the two BZLF1 responsive elements ZRE 3 and ZRE 4 in the promoter of *BBLF4* as in Fig. 3A. **(B)** EMSA results of BZLF1 and the DNA templates ZRE 3+4, ZRE 0+4 and ZRE 3+0 suggested a cooperative binding of BZLF1 to ZRE 3 and ZRE 4. The ZRE 3+4 template was robustly bound by BZLF1, which interacted less efficiently with ZRE 3+0 and barely with ZRE 0+4. **(C)** Individual Hill slope curves of BZLF1 binding to the mononucleosomal DNA templates ZRE 3+4, ZRE 0+4 and ZRE 3+0 show the result of three independent experiments. **(D)** The position of the ZRE 3 motif within the DNA template ZRE 3+0 was altered as illustrated in (A). BZLF1 was competent to bind its ZRE 3 site in two more proximal positions (ZRE +15 nt and +30 nt) within the nucleosomal core.

**Fig. 5.**
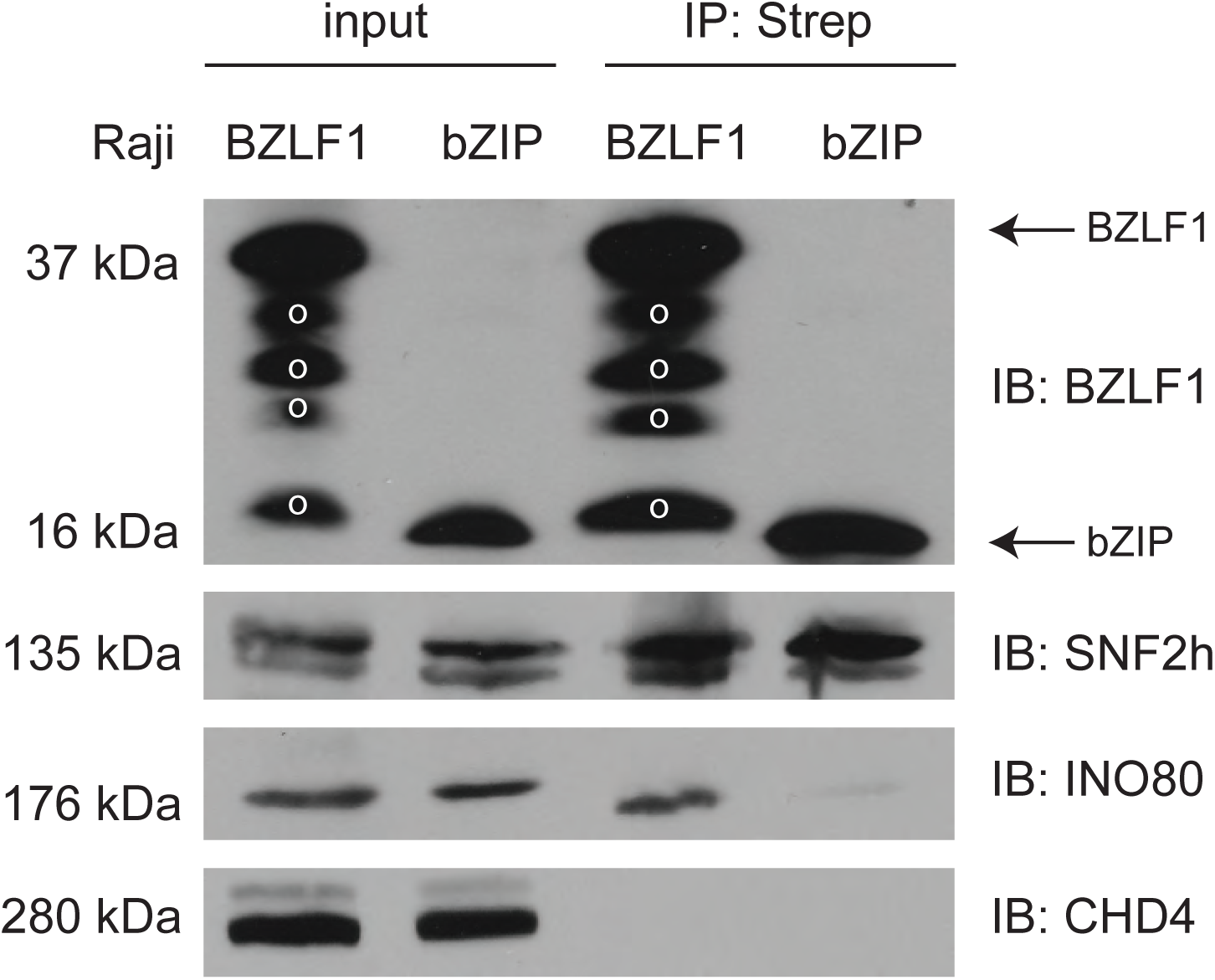
BZLF1 interacts with the core subunits of the cellular chromatin remodelers SNF2h and INO80 *in vivo*. Raji cell lines were stably transfected with tetracycline regulated expression plasmids encoding Strep-tagged BZLF1 full-length or bZIP protein (Fig. S1) consisting of aa175 to aa236 with the DNA binding and dimerization domains of BZLF1. After treatment with benzonase and DNase I, the Strep-tag fusion proteins of lytically induced cells were captured with Streptavidin beads. Co-precipitated proteins were analyzed with antibodies directed against SNF2h, INO80, and CHD4. In the immunoblot detecting BZLF1 and bZIP (top panel), 0.5 % of the total protein lysate was loaded as “input” per lane; in each of the two lanes labeled “IP:Strep” 10 % of the immunoprecipitated material was loaded. In the immunoblots detecting SNF2h, INO80, or CHD4, 1 % of the total protein lysate was loaded as “input” per lane; in the lanes labeled “IP:Strep” 90 % of the immunoprecipitated material was loaded per lane. ‘o’ indicates signals from proteolytic degradation.

**Fig. 6.**
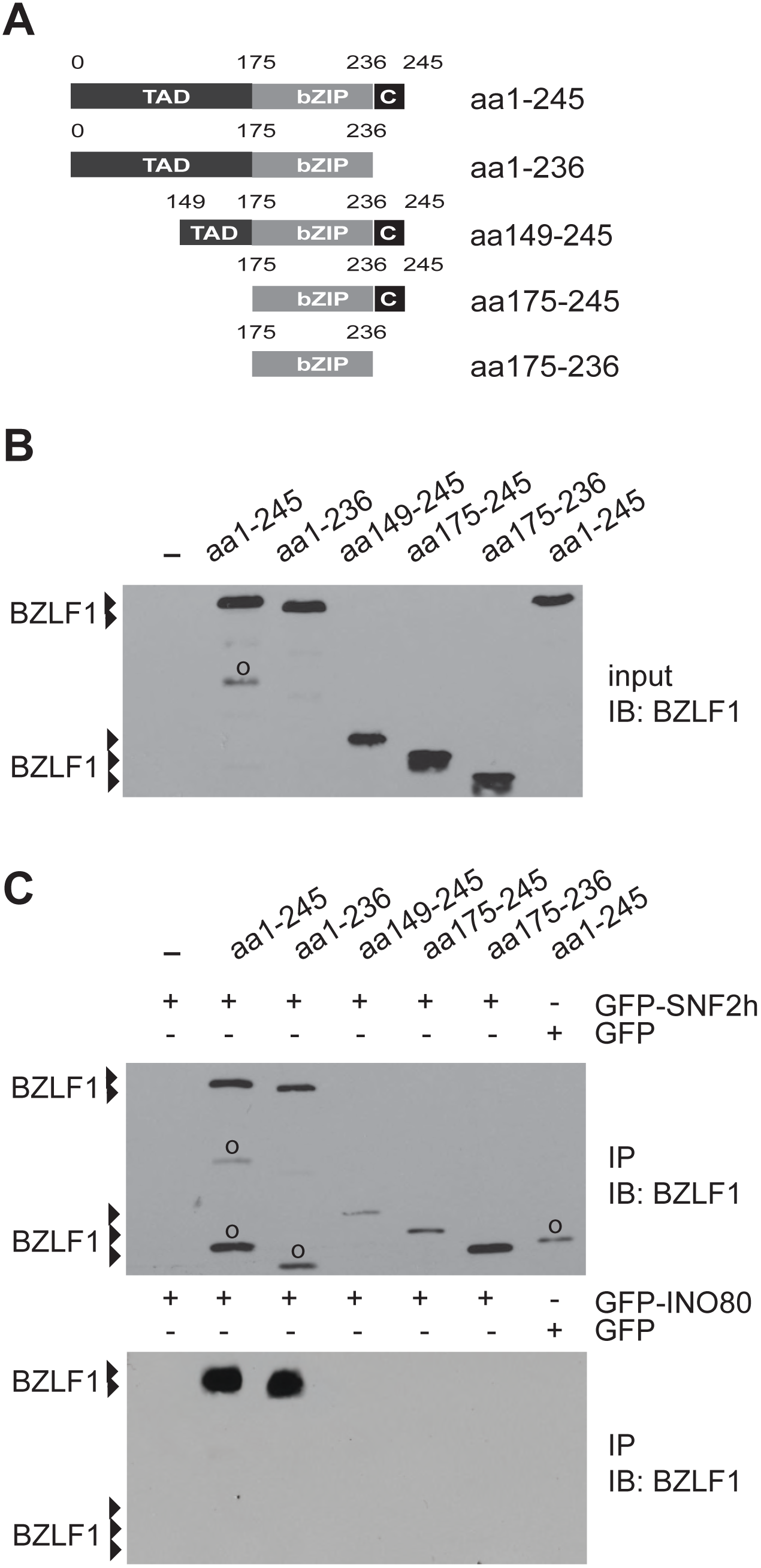
BZLF1’s transactivation and bZIP domains promote protein-protein interactions with chromatin remodelers. **(A)** Shown are the modular structures of truncated BZLF1 variants. **(B)** Protein expression of the truncated BZLF1 variants (A) in HEK293 cells was analyzed by immuno detection with the BZLF1-specific BZ1 antibody. **(C)** HEK293 cells were co-transfected with expression plasmids encoding BZLF1 variants and GFP-tagged chromatin remodeler ATPase subunits SNF2h or INO80. Cell lysates were treated with the enzymes benzonase and DNase I prior to immunoprecipitations of the GFP-tagged chromatin remodelers with GFP-binder beads. The analysis of input and immunoprecipitated material was done by Western blot detection with the BZ1 antibody directed against BZLF1. ‘o’ indicates signals from proteolytic degradation.

**Fig. 7.**
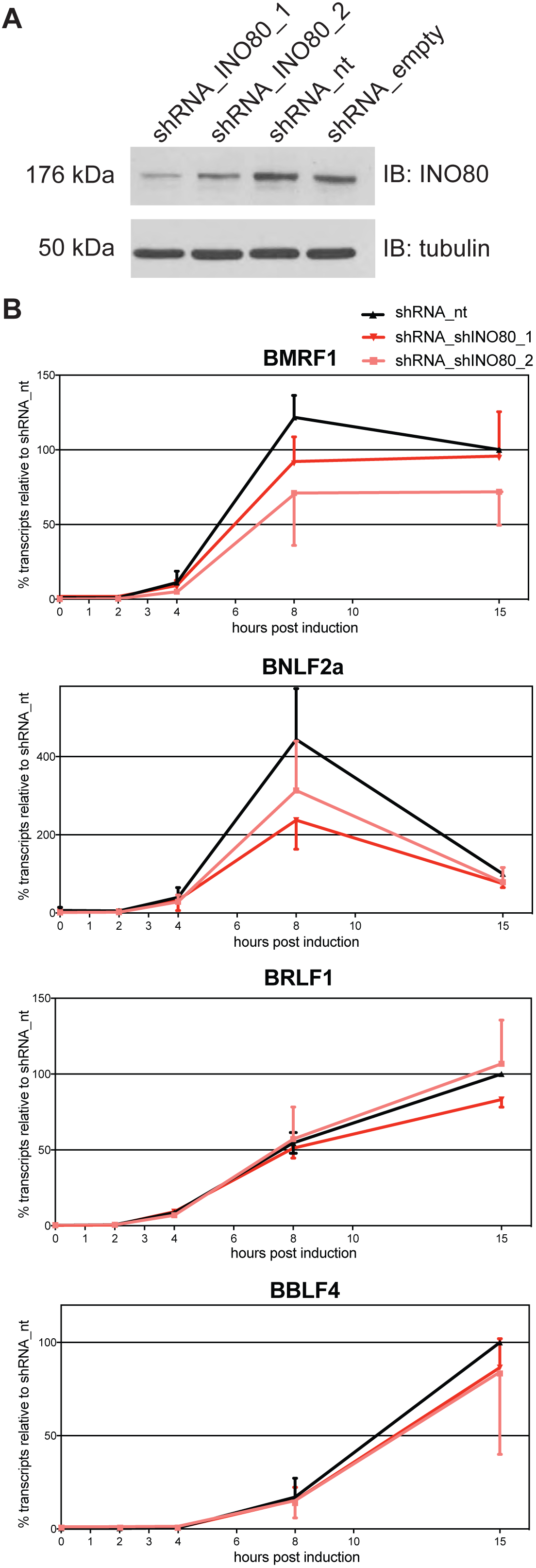
shRNA knock-downs of INO80 reduce the rapid lytic transcriptional reactivation of certain early lytic EBV genes. **(A)** Knock-down efficiency of INO80 protein in Raji p4816 cell lines transduced with lentiviral vectors encoding shRNA_INO80_1 and shRNA_INO80_2 directed against INO80 transcripts or shRNA_nt (non-targeting) and shRNA_empty (no shRNA) as controls. Tubulin served as a loading control. One representative experiment out of three is shown. **(B)** Transcripts of selected viral early lytic genes (BMRF1, BNLF2a, BRLF1, BBLF4) in Raji p4816 cell lines transduced with different shRNAs (shRNA_INO80_1, shRNA INO80_2, or shRNA_nt as a negative control) were quantified by RT-qPCR after different time points post induction with 100 ng/ml doxycycline as indicated (2, 4, 8, or 15h or non-induced, i.e. 0h). Values obtained with the Raji p4816 cell line transduced with the control shRNA_nt at 15h post induction were set to 100 %. Mean values and standard deviations from three independent experiments are shown. Primers used in RT-qPCR reactions are shown in Table S2 and individual experiments are shown in Figure S10.

### Absolute quantification of transcripts using RT-qPCR

Raji p4816 cells which express lentivirally transduced shRNA_nt or two INO80 targeting shRNAs (shRNA_INO80_1 and _2) were induced for BZLF1 expression (100 ng/ml doxycycline) for 2, 4, 8 or 15 hours. From non-induced and from induced cells fractions were collected at the indicated time points to determine the induced expression of GFP by flow cytometry and to analyze the INO80 protein levels by immunoblot detection. At each time point 1×10^6^ cells were centrifuged, washed with ice cold PBS, and pellets were snap frozen in −80 °C. Isolation of cellular RNA was accomplished with the RNeasy Mini Kit (Qiagen). The cell lysates were homogenized with QiaShredder columns (Qiagen). The subsequent steps were performed according to the manufacturer’s instructions. RNA was eluted in 30 μl of RNase-free water. Contaminating genomic DNAs were removed with RNAse-free deoxyribonuclease I (DNase I, Thermo Fisher Scientific) in the presence of RNase inhibitor (Roche) at room temperature for 40 min followed by inactivation at 65 °C for 10 min. The efficiency of DNA digestion was determined by PCR amplification of viral DNA with the BBLF2 primer pair (Tab. S2). RNA was reverse-transcribed into cDNA using the SuperScript III First Strand Synthesis SuperMix Kit (Thermo Fisher Scientific) following the manufacturer’s instructions. cDNA was analyzed quantitatively by RT-qPCR using the LightCycler480 (Roche) instrument.

### Statistical analysis

We used Prism 6 (GraphPad) for statistical analysis and the two-tailed ratio T-test was applied unless otherwise mentioned.

## RESULTS

### Loss of histone H3 at repressed lytic promoters follows initial lytic viral reactivation

We used our established model (Woellmer et al., 2012) to analyze the kinetics of nucleosomal loss at selected loci in EBV DNA upon lytic induction. Raji cells, a Burkitt’s lymphoma derived cell line latently infected with EBV, were engineered to contain an inducible BZLF1 allele and termed Raji p4816 cells (Fig. S1). In this model, adding doxycycline triggers the expression of BZLF1 and induces viral lytic reactivation. We compared the kinetics of BZLF1’s induced expression with the kinetics of nucleosome loss at selected, BZLF1-controlled lytic EBV promoters. We harvested samples from uninduced and induced Raji p4816 cells at different time points after addition of 100 ng/ml doxycycline and analyzed BZLF1 expression by Western blotting (Fig. 1A). Raji p4816 cells showed a clear BZLF1 signal already two hours post induction, and the protein level increased and reached high levels 15 hours post induction. As expected, doxycycline did not induce BZLF1 in parental Raji cells (Fig. 1A).

We were concerned if the level of BZLF1 protein present in induced Raji p4816 cells might exceed the levels present in EBV-positive cells that support EBV’s lytic phase. To address this point, we compared the levels of BZLF1 in the B95-8 cell line with levels in our Raji cell model after induced expression of BZLF1. A small fraction of B95-8 cells spontaneously enter the lytic phase and support virus *de novo* synthesis (Miller et al., 1972). We found that the BZLF1 levels we reach in the Raji inducible system are in a range also found in the small fraction of B95-8 cells that undergo the lytic cycle of EBV (unpublished data).

Next, we performed chromatin immunoprecipitation (ChIP) with cross-linked viral chromatin, which had been fragmented to an average size of 150 base pairs (bp) and an antibody directed against histone H3 indicative of the histone octamer. We detected a partial loss of H3 at promoter sites of certain early lytic genes as reported previously (Woellmer et al., 2012), but only after 15 hours post induction (Fig. 1B). In contrast, H3 levels were unaffected at latent and late lytic promoters (Fig. 1B) (Woellmer et al., 2012). The data indicated that BZLF1 expression clearly preceded the detectable loss of H3 at certain promoters of early lytic genes in lytically induced Raji cells.

Addition of doxycycline to mammalian cells might have adverse effects and alter transcription and chromatin structure or affect cell vitality. We tested this aspect in parental Raji cells and at doxycycline concentrations used in this and all subsequent experiments. RNA-Seq experiments with induced and uninduced Raji cells did not identify any noticeable change in cellular transcription (unpublished data).

### BZLF1 binds mononucleosomal DNA in viral lytic promoters *in vivo*

It is unclear if BZLF1 can bind nucleosomal DNA directly or relies on a mechanism that exposes ZRE motifs, presumably by nucleosome eviction prior to BZLF1’s binding. The former possibility would correspond to pioneer factor-like binding of BZLF1 at nucleosomal sites; the latter would involve additional unknown factors required to facilitate BZLF1’s binding. To examine the first possibility, we looked for co-occupancy of BZLF1 and histone octamers at ZREs in our model cell line *in vivo*. We performed ChIP and sequential ChIP (ReChIP) experiments with latent phase chromatin at different time points after lytic induction with antibodies directed against BZLF1 and the histone mark H3K4me1. Prior to these ChIP experiments, the chromatin had been crosslinked and sheared to mononucleosomal size by sonication. We chose an antibody directed against the specific H3K4me1 histone mark for two reasons: (i) in our hands, antibodies directed against pan H3 (and other core histone proteins tested) performed adequately in classical ChIP experiments, but poorly in ReChIP experiments. In contrast, antibodies directed against certain histone marks such as H3K4me1 were well suited for the technically challenging ReChIP experiments. (ii) Upon induction of EBV’s lytic phase, the prevalence of the H3K4me1 modification increased slightly at early lytic promoters over time (to be published and Fig. 2A), which improves the chances to detect possible interactions of BZLF1 with ZREs embedded in nucleosomal DNA in ReChIP experiments.

BZLF1’s binding (Fig. 2A, left panel) was detected at most early lytic promoters within four hours after lytic induction. A modest, up to two-fold increase of H3K4me1 (right panel) became obvious seven hours post induction. Results from individual experiments are shown in Fig. S2. ReChIP experiments were done with either order of the two antibodies (Fig. 2B). The results demonstrated the co-occupancy of BZLF1 and histone octamers on the same DNA molecules in the promoter regions of early lytic genes seven and fifteen hours post induction. ReChIP experiments in which a non-specific IgG antibody replaced either of the two antibodies served as negative controls for the second precipitation step (Fig. 2C). Carry-over of chromatin complexes from the first round of ChIP experiments with antibodies directed against BZLF1 or H3K4me1 was low (Fig. 2C, left panel) or negligible (Fig. 2C, right panel), respectively. The results suggested that BZLF1 can bind directly to nucleosomal DNA *in vivo*.

### BZLF1 binds mononucleosomal DNA *in vitro*

We verified our *in vivo* finding in a defined and unambiguous *in vitro* system using electromobility shift assays (EMSAs) with a purified BZLF1 protein and ZRE-containing DNA fragments in their free states or bound as mononucleosomes (Fig. 3). The latter bound fragments serve as surrogates for viral chromatin in its repressed state.

The promoter of the early lytic gene BBLF4, which encodes the viral DNA helicase, harbors five ZREs of nine base pairs in length. All five ZREs contain CpG dinucleotides and show a methylation-dependent binding of BZLF1 (Bergbauer et al., 2010). We prepared three 156 bp long DNA fragments derived from this promoter region that differed in the positioning of two ZREs, ZRE 3 and ZRE 4 (Fig. 3A). A 156 bp fragment of the BRLF1 coding sequence, which lacks ZREs and is not bound by BZLF1, served as negative control. All four 156 bp fragments, which had been fully CpG methylated using a commercially available *de novo* CpG methyl transferase were reconstituted into mononucleosomes by salt gradient dialysis using *Drosophila* embryo histone octamers (Krietenstein et al., 2012) (Fig. S3A, B). Strep-tagged BZLF1 protein was expressed in HEK293 cells and purified under native conditions by Strep-Tactin affinity chromatography (Fig. S3C, D).

EMSAs with these purified reagents (Fig. 3B) allowed measuring BZLF1’s binding to the four DNA fragments in their free (upper row of panels) or mononucleosomal (lower row of panels) states and the determination of the respective equilibrium dissociation constants (K_D_) (Fig. 3C). BZLF1 bound with similar affinity (K_D_ of approx. 10 to 20 nM) to the three ZRE-containing and histone-free DNA fragments consistent with previous experiments (Bergbauer et al., 2010) and independent of the number of ZREs. BZLF1 bound only weakly to the control fragment lacking a ZRE (Fig. 3B), which is consistent with the widely observed, low-level but non-specific DNA binding of transcription factors (Fried and Crothers, 1981). BZLF1’s binding to free DNA resulted in several shifted bands, which is a common observation (Bergbauer et al., 2010).

In contrast, BZLF1’s binding to mononucleosomal DNA differed for the three ZRE-containing fragments (Fig. 3B, lower row of panels). BZLF1 did not bind to mononucleosomes without (control fragment) or with only one ZRE (ZRE 3 or ZRE 4). The well-studied yeast transactivator Pho4, which served as a negative control because it does not bind to nucleosomal sites (Venter et al., 1994), did not yield shifted bands with its binding site buried in mononucleosomes but did with free DNA (Fig. S4). However, BZLF1 did bind to the ZRE 3+4 fragment, again with a K_D_ of about 13 nM (Fig. 3C). A truncated BZLF1 protein (aa149-245) that lacks the activation but retains the DNA-binding domain, bound both free and mononucleosomal ZRE 3+4 fragments (Fig. S5) indicating that the DNA-binding domain is sufficient to mediate the pioneer factor-like binding of BZLF1.

The binding of BZLF1 to the mononucleosomal ZRE 3+4 fragment (but not to fragments with single ZRE 3 or ZRE 4 motifs) suggested that BZLF1 requires at least two binding sites for stable binding to a nucleosome. As an alternative interpretation, two ZREs might be required to outcompete the histone octamer for binding such that the shifted complex migrated like a complex of BZLF1 with free DNA. We ruled out this latter possibility by comparing BZLF1 complexes with free and mononucleosomal ZRE 3+4 fragments run in parallel in the same gels (Fig. 3B, bottom row, rightmost panel, and Fig. 3D). The migration position of BZLF1 in complex with free DNA differed from that in complex with mononucleosomal DNA. An anti-FLAG antibody appropriate for binding FLAG-tagged BZLF1 supershifted the signals and unambiguously identified BZLF1 in both the free and the mononucleosomal DNA shift complexes (Fig. 3D).

Yet another interpretation could be that BZLF1 did not necessarily require two ZREs for binding on a nucleosome, but that a ZRE had to be close to the entry and or exit points of the nucleosomal DNA rather than close to the dyad. It is known that nucleosomal DNA can undergo thermal “breathing” motions that transiently expose DNA sites close to the exit/entry points but much less frequently sites close to the dyad (Anderson and Widom, 2000). To investigate this possibility, we modified the ZRE 3 or ZRE 4 sequences in the ZRE 3+4 fragment by PCR mutagenesis such that we obtained two fragments with only one ZRE located at different positions relative to the entry/exit points termed ZRE 0+4 and ZRE 3+0 (Fig. 4A). EMSAs with BZLF1 and these two constructs in mononucleosomal forms demonstrated that BZLF1’s binding to ZRE 0+4 was barely detectable (Fig. 4B, middle panel), relatively strong to ZRE 3+0 (Fig. 4B, right panel), and strongest to ZRE 3+4 (Fig. 4B, left panel). From Hill plots (Fig. 4C) we found again a dissociation constant of 13 nM for BZLF1 binding to the ZRE 3+4 mononucleosome compared to a K_D_ of about 100 nM for ZRE 3+0. The dissociation constant could not be determined for ZRE 0+4 mononucleosomes. In contrast and as expected, the K_D_ values of BZLF1 binding to the different free DNA fragments were in the range of 10 to 20 nM in three independent experiments (Fig. S6).

We made use of the clearly detectable binding of BZLF1 to the single ZRE in the ZRE 3+0 fragment to ask if thermal “breathing” or nucleosomal phasing played a major role to this binding. To do so we altered the position of the single ZRE in the ZRE 3+0 fragment relative to the original position of this ZRE by −5 nucleotides (nt), +10 nt, +15 nt, and +30 nt as shown in Fig. 4A. With these four constructs, we repeated the EMSA analysis and observed robust BZLF1 binding to three of four mononucleosomes tested (−5 nt, +15 nt, and +30 nt), in all cases stronger than to the ZRE 0+4 mononucleosome (Fig. 4D vs. B). BZLF1 binding to ZRE 3+0 +10 nt (Fig. 4A) was not detectable (Fig. 4D, middle panel). This finding discredits a prominent role of DNA “breathing” for BZLF1’s binding, especially given that the ZRE is more internal in the ZRE 3+0 +30 nt than in the ZRE 0+4 fragment. The fact that BZLF1 did not bind to ZRE 3+0 +10 nt (Fig. 4D, middle panel) is reminiscent of ZRE 0+4 (Fig. 4B, middle panel), because both binding sites are positioned similarly, i.e. 18 and 17 nt from the distal ends (Fig. 4A).

The data do not seem to support nucleosomal phasing as a critical determinant either. Shifting of the single ZRE in the fragment ZRE 3+0 by multiples of 15 bp (+15 nt versus +30 nt), which corresponds roughly to one and a half helical turns on the histone octamer surface (44), did not affect BZLF1 binding (Fig. 4D) suggesting that it did not matter which part of the ZRE faces inward or outward from the nucleosome. Taken together, we conclude that the properties and the position of a given ZRE (ZRE 3 versus ZRE 4) and, to a much larger degree, the cooperation between two ZREs (ZRE 3+4 construct) support BZLF1’s binding to a nucleosomal site, but the data do not provide an obvious rule of BZLF1 binding to nucleosomal DNA.

### BZLF1 and cellular chromatin remodeling enzymes interact

In our *in vitro* experiments with reconstituted nucleosomes we did not observe a histone loss or a disassembly of the nucleosome upon BZLF1’s binding because shifted bands characteristic of a BZLF1-free DNA complex (Fig. 3D) or an increase of free DNA (Fig. 3B, bottom row, rightmost panel) were not detected. It thus appeared that BZLF1’s binding to chromatin and the ejection of nucleosomes *in vivo* are two distinct but possibly linked processes.

We hypothesized that *in vivo* BZLF1 might first bind at ZREs in promoter elements of lytic target genes on top of the nucleosomes without ejecting them, but then recruit cellular chromatin remodeling enzymes that would mediate the loss of histones. With a candidate approach we analyzed whether BZLF1 interacted with members of the chromatin remodeler families, ISWI and INO80. We co-immunoprecipitated BZLF1 and the catalytic ATPase subunits SNF2h (encoded by the gene *SMARCA5* and corresponding to the paradigmatic *Drosophila* ISWI ATPase (Flaus et al., 2006)) or INO80 (Shen et al., 2000). The chromatin remodelers were both expressed at endogenous levels in Raji cells, while BZLF1 was expressed in cells stably transfected with our doxycycline inducible expression system (Fig. S1) encoding Strep-tagged BZLF1 full-length protein (aa1-245) or only the BZLF1 bZIP domain (aa175-236). After over-night induction, cell lysates were treated with benzonase and DNase I prior to immunoprecipitation to eliminate nucleic acid-mediated recovery of the factors. Tagged BZLF1 was immunoprecipitated on Streptavidin beads and co-precipitation of chromatin remodelers was determined by subsequent Western blotting (Fig. 5). This approach revealed interactions of full-length BZLF1 with endogenous SNF2h and INO80 protein, but not with CHD4. The BZLF1 bZIP domain alone, i.e. BZLF1 lacking its transactivation domain (TAD) and the ultimate carboxy-terminus, interacted with SNF2h but not with INO80 (Fig. 5). These results suggested that BZLF1 interacts with subunits of at least two cellular chromatin remodeler families possibly to recruit them to lytic gene promoters and support EBV’s lytic reactivation.

### Different BZLF1 domains mediate the interaction with SNF2h versus INO80

The differential binding of the bZIP constructs to SNF2h versus INO80 indicated that different domains of BZLF1 might mediate these interactions. We extended our co-immunoprecipitations to include more BZLF1 derivatives (Fig. 6A). We transiently transfected HEK293 cells with the BZLF1 constructs, which were co-expressed with the GFP-tagged chromatin remodeler ATPase subunits, SNF2h or INO80. The BZLF1 expression plasmids were adjusted to obtain similar protein levels (Fig. 6B). Cell lysates were again treated with benzonase and DNase I, immunoprecipitations were done with GFP-binder coupled sepharose beads and the co-precipitated BZLF1 was detected by Western blotting using the antibody directed against a motif within BZLF1’s bZIP domain. The immunoblots (Fig. 6C; controls in Fig. S7) demonstrated that SNF2h interacted with all BZLF1 derivatives tested (upper panel) probably identifying aa175-236 of BZLF1 as the domain responsible for SNF2h interaction. In contrast, the intact transactivation domain of BZLF1 was essential to bind INO80 or components of the INO80 remodeler complex (Figs. 6C and S7), a feature, which is in agreement with the presumed functions of activation domains of DNA-binding transcription factors in general. Therefore, we put our focus on INO80 in most of the following experiments, in which we investigated the functional role of chromatin remodelers in lytic viral reactivation.

### BZLF1 supports the recruitment of the INO80 chromatin remodeler complex to viral DNA

The transactivating domain of BZLF1 appeared to interact with INO80 or components of the INO80 remodeler complex. We wondered whether BZLF1 might as well recruit this chromatin remodeler to viral chromatin, where it could induce local histone loss as seen in Fig. 1, for example. It is technically very challenging to perform convincing ChIPs with remodeler complexes in yeast, drosophila, or mammalian cells probably because these multicomponent complexes are large and presumably make only transient contacts with DNA binding factors and chromatin. We followed the experimental work by the Tsukiyama lab (Gelbart et al., 2005), who used a catalytically inactive remodeler mutant to map its targets genome-wide. We engineered an HA-tagged INO80 allele with a point mutation (W653Q), which makes this remodeler dysfunctional. An expression plasmid encoding the mutant INO80 protein was transiently transfected alone or in conjunction with a plasmid encoding BZLF1 into 2089 EBV HEK293 cells that are latently infected with the recombinant EBV strain B95-8 (Delecluse et al., 1998). Upon BZLF1 expression, these cells readily support the induction of EBV’s lytic phase and the expression of all lytic genes (Delecluse et al., 1998). Fifteen hours after DNA transfection, ChIPs were performed with an HA-tag specific antibody. The results suggested that the level of HA-tagged INO80 (W653Q) is increased as a function of co-expressed BZLF1 at selected viral promoters (Fig. S8). In three independent experiments the increase was modest but reproducible and is in line with the notion that BZLF1 recruits INO80 to these viral promoters. This effect can also be seen to some degree at the control locus (*HPRT1*) and a latent viral gene (EBER) (Fig. S8), which might indicate a global alteration of chromatin structure upon lytic phase induction, because BZLF1 binds to more than 10^5^ binding sites in cellular chromatin (unpublished data).

### shRNA mediated knock-down of INO80 reduces transcriptional reactivation of certain early lytic genes of EBV

Next, we asked whether INO80 levels might be critical to activate viral lytic genes upon expression of BZLF1. We engineered lentiviruses to stably express shRNAs directed against INO80 transcripts in Raji p4816 cells (Fig. S9A, B) and tested the timely expression of selected viral lytic genes upon addition of doxycycline by RT-qPCR.

As can be seen in Fig. 7A, two shRNAs efficiently reduced the steady-state levels of INO80 protein in Raji cells. We next analyzed the transcriptional activation of four early viral genes (BMRF1, BNLF2a, BRLF1, BBLF4) in three different Raji p4816 cell lines stably transduced with shRNA_INO80_1, shRNA_INO80 _2, or a non-targeting shRNA_nt control (Fig. S9A, B). The knock-down of INO80 resulted in a reduced activation of BMRF1 and BNLF2a eight hours post induction (Fig. 7B). In contrast to these two ‘early responding’ genes, BRLF1 and BBLF4, which have considerably slower kinetics of induction, showed a very modest reduction of their transcript levels 15 hours post induction, only (Fig. 7B and Fig. S10). We concluded from this experiment that INO80 may play a critical functional role in the early phase of viral reactivation at certain viral promoters of early lytic genes.

### siRNA-mediated knock-down of INO80 inhibits *de novo* synthesis of virus

As BZLF1 is the crucial trigger for viral reactivation and interacts with at least two cellular chromatin remodelers, we hypothesized that at least one of them should be necessary for lytic induction. To test this hypothesis, we used an siRNA knock-down strategy to assess the roles of SNF2h or INO80 in 2089 EBV HEK293 cells (Delecluse et al., 1998). Upon transient transfection of a BZLF1 expression plasmid (Hammerschmidt and Sugden, 1988), this 2089 EBV HEK293 producer cell line releases infectious virus, which can be quantified by assaying infected, GFP-positive Raji cells by flow cytometry (Steinbrück et al., 2015).

The 2089 EBV HEK293 cells were treated for three days with siRNA pools targeting the *SMARCA5* gene encoding SNF2h or the *INO80* gene or with a non-targeting control siRNA. The respective knock-down efficiencies were assessed by Western blotting (Fig. 8A). In these siRNA-treated cells virus synthesis was initiated by transient co-transfection of expression plasmids encoding BZLF1 together with gp110/BALF4 as described (Neuhierl et al., 2002). Expression of gp110/BALF4 increases virus infectivity by about a factor of ten (Neuhierl et al., 2002). Three days after plasmid co-transfection, cell supernatants were collected and defined volumes were used to infect Raji cells. The fractions of GFP-expressing Raji cells were determined by flow cytometry after three additional days such that the virus concentrations could be calculated (Fig. 8B).

**Fig. 8.**
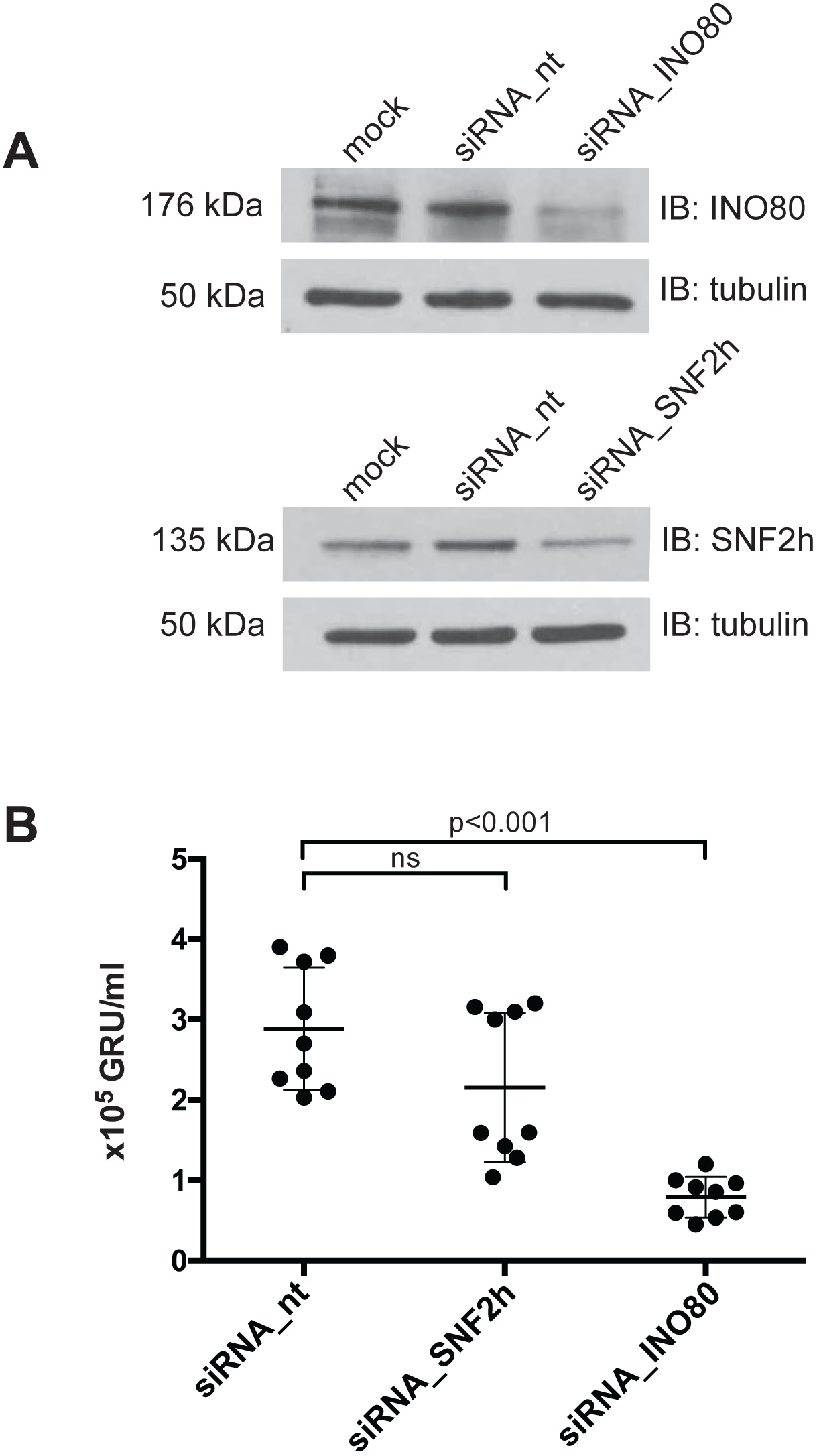
siRNA knock-down of SNF2h and INO80 subunits reduce virus *de novo* synthesis. **(A)** Knock-down efficiencies of the SNF2h and INO80 ATPase subunits in 2089 EBV HEK293 cells were analyzed by immunoblotting. Tubulin served as a loading control. One representative experiment out of three is shown. **(B)** Quantification of virus concentrations released after lytic induction of 2089 EBV HEK293 cells indicate a reduction of virus synthesis after INO80 knock-down. Mean and standard deviations from nine independent experiments are shown. ns=not significant. p values of an unpaired t-test corrected with the Sidak-Bonferroni method are shown. GRU, green Raji units.

Steady-state protein levels of SNF2h or INO80 were modestly reduced after three days of siRNA treatment (Fig. 8A). Cells treated with non-targeting siRNAs or with an SNF2h-specific siRNA pool (Fig. S9C) did not differ significantly in the levels of released, infectious EBV (Fig. 8B). However, cells treated with an INO80-specific siRNA pool released significantly fewer viral particles (Fig. 8B). This observation is notable given the only modest diminution of INO80 by the siRNA treatment (Fig. 8A). None of the siRNA pools directed against SNF2h or INO80 had an adverse effect on cell viability (Fig. S9D) suggesting that the reduced virus synthesis after siRNA knockdown is a specific and INO80-related effect.

Together, these results support a critical role for INO80 and for BZLF1’s acting as a pioneer factor in EBV lytic activation.

### Discussion

EBV takes advantage of the host cell’s epigenetic machinery to establish a stable latent infection. Upon infection, the viral DNA is epigenetically naïve, i.e. free of histones and devoid of methylated CpG dinucleotides (Kalla et al., 2010; Kintner and Sugden, 1981; Fernandez et al., 2009). In the course of establishing the latent phase, the host cell’s epigenetic machinery compacts the viral DNA into nucleosomal arrays, introduces repressive histone modifications, and initiates the methylation of the majority of viral CpG dinucleotides (Kalla et al., 2010). As a consequence, viral promoters, with the exception of those of a few latent genes, are silenced during EBV’s latent phase. Densely positioned nucleosomes, repressive histone marks introduced by Polycomb proteins, and extensive DNA methylation keep the virus in a strictly latent, dormant mode (Ramasubramanyan et al., 2012; Woellmer et al., 2012).

EBV can escape from latency and enter the lytic, productive phase when its host B cells terminally differentiate to plasma cells (Laichalk and Thorley-Lawson, 2005). In the lytic phase, the loss of nucleosomes increases the accessibility of viral DNA to binding transcription factors. The removal of repressive and the gain of active histone marks reactivate the promoter regions of early viral lytic genes, enabling the virus to replicate its DNA, express late viral genes, and produce viral progeny to infect new B cells (Hammerschmidt, 2015 for a recent review).

The viral transcription factor BZLF1, which is induced upon terminal plasma cell differentiation, is the switch triggering a transition from the latent to the lytic phase. First, BZLF1 induces the expression of several early lytic EBV genes by binding sequence-specifically to ZREs in their promoter regions. Many ZREs need to contain 5-methyl cytosine residues to permit BZLF1’s binding and thus CpG methylation of viral DNA is a prerequisite to express certain, essential, early lytic genes (Kalla et al., 2010; Kalla et al., 2012). Second, BZLF1 enables viral DNA replication. It binds to the lytic origin of DNA replication and promotes the recruitment of components of the viral DNA replication machinery to this viral origin to support its function (Schepers et al., 1993). Third, BZLF1 directly or indirectly causes the loss of nucleosomes in the promoter regions of viral lytic genes, which correlates with their expression (Hammerschmidt, 2015).

We and others have hypothesized that the efficient reactivation of silenced, inactive viral chromatin is forced by a presumed pioneer function of the BZLF1 transcription factor (Woellmer et al., 2012), chromatin alterations (Adamson and Kenney, 1999; Zerby et al., 1999), and/or the additional recruitment of chromatin remodelers (Woellmer et al., 2012). Our results, both *in vivo* (Fig. 2) and *in vitro* (Figs. 3 and 4), show that BZLF1 can bind mononucleosomal DNA in promoter regions of early lytic genes known to be regulated by BZLF1. We have also shown that BZLF1 can bind ZREs in nucleosomes close to the nucleosome dyad (Fig. 4D). The nucleosome-binding activity is encoded within the C-terminal part of BZLF1 encompassing the bZIP domain and C-terminus and does not depend on BZLF1’s transactivation domain (Fig. S5). BZLF1’s binding does not eject nucleosomes *in vitro* (Fig. 3B, D) in the absence of other molecular machines. This observation is in line with our initial data (Fig. 1B) indicating that the binding of BZLF1 precedes a decrease in nucleosomal occupancy at early lytic promoters by hours.

Transcriptional activation of lytic viral genes (BMRF1 and BNLF2a in Fig. 7B for example) appears to precede nucleosome ejection as shown in Fig. 1B (15 h post induction). This lag is probably due to the fact that the Raji cell line contains about 15 copies of viral genomes per cell. The loss of histones can be detected, only, when a considerable percentage of nucleosomes are removed on the background of the many EBV genome copies per cell. It is unlikely that they all become activated and respond synchronously to the expression of BZLF1. The knockdown of INO80 reduces transcriptional activation of BMRF1 and BNLF2a genes 8 hours post induction (Fig. 7B) but not later. This observation supports our working hypothesis and suggests that BZLF1 binds to nucleosomal DNA and concomitantly recruits chromatin remodelers such as INO80 and the transcriptional machinery, which together open viral chromatin, revert epigenetic silencing and promote gene activation.

Co-immunoprecipitations suggest the *in vivo* interactions of BZLF1 with the ATPase subunits SNF2h and INO80 of the chromatin remodeler families ISWI and INO80, respectively (Fig. 5). In Herpes Simplex Virus 1 (HSV-1) VP16, BZLF1’s functional counterpart, regulates the lytic reactivation process. VP16’s transactivation domain recruits general transcription factors and chromatin cofactors, including chromatin remodeling enzymes to sites in viral promoters, dramatically reducing their histone occupancy (Neely et al., 1999; Herrera and Triezenberg, 2004). The ATPase subunit SNF2h has been reported previously to promote HSV-1 immediate-early gene expression as well as replication and might also interact with VP16 (Bryant et al., 2011).

Similarly, the INO80 remodelers are able to slide nucleosomes, exchange histones, regulate transcription, and are involved in DNA repair and cell cycle checkpoint control (Shen et al., 2000; Tsukuda et al., 2005; van Attikum et al., 2007). While INO80 has not been implicated in the reactivation of herpes viruses, the EBV nuclear proteins EBNA-LP and EBNA2, which are necessary for lymphoblastoid cell line growth and survival have been found associated with an INO80 remodeler complex (Portal et al., 2013). Our results indicate that BZLF1’s transactivation domain interacts with the INO80 ATPase subunit (Fig. 6C) and likely recruits the entire remodeler complex to viral chromatin, where we envision that it mobilizes histone octamers and disrupts the nucleosome-dense regions of the viral promoters upon lytic reactivation. INO80 knock-down experiments support this critical role (Fig. 7B).

Recruitment of INO80 via BZLF1’s transactivation domain is consistent with our identification of a subgroup of ZREs in viral DNA with a higher than average nucleosome occupancy during latency. Nucleosome loss and the formation of hypersensitive sites at these ZRE elements was only observed with full-length BZLF1 protein but not with the bZIP domain lacking BZLF1’s transactivation domain (Fig. 2C, bottom panel, in reference Woellmer et al., 2012). In line with this observation, the interaction of BZLF1 and INO80 also depended on the intact transactivation domain of BZLF1 (Figs. 5 and 6). All our observations support the notion that BZLF1 recruits the INO80 chromatin remodeling complex to lytic promoters in viral chromatin and orchestrates their loss of nucleosomes.

Recent reports support the view that pioneer transcription factors can penetrate epigenetically silenced chromatin to recognize and bind their DNA-binding sites activating the associated genes. The cell type-specific distribution of histone marks plays an important role in the recruitment of pioneer factors, as some are capable of reading them. For example, the recruitment of FoxA to enhancers depends on epigenetic changes of enhancer hallmarks (Sérandour et al., 2011). The nucleosome-binding activity of FoxA is facilitated by the histone marks H3K4me1 and H3K4me2 but not by histone acetylation (Cirillo and Zaret, 1999; Lupien et al., 2008). FoxA can even favor H3K4me2 deposition (Smale, 2010). The pioneer factor PBX1 is also capable of reading specific epigenetic signatures such as H3K4me2 (Magnani et al., 2011a; Berkes et al., 2004), and PU.1 reprograms the chromatin landscape through the induced deposition of H3K4me1 (Heinz et al., 2010). The viral transcription factor BZLF1 might read and decipher epigenetic modifications as well. In fact, we confirmed in ReChIP experiments that BZLF1 co-occupies H3K4me1 in the viral promoters of early lytic target genes (Fig. 2B).

The viral transcription factor BZLF1 belongs to the basic leucine-zipper (bZIP) family of transcription factors and contains a variant of the leucine zipper motif responsible for the coiled-coil structure (Farrell et al., 1989; Chang et al., 1990; Lieberman and Berk, 1990). BZLF1 forms homodimers and binds DNA motifs via its two long bZIP helices (Petosa et al., 2006). The basic region of each helix contacts the major groove and the zipper region forms a coiled-coil stabilized by BZLF1’s C-terminal tail. The closest relative of BZLF1 is the cellular c-Fos/c-Jun heterodimer AP1 transcription factor (Farrell et al., 1989). Its binding to nucleosomal DNA is reduced compared to free DNA sequence motifs, but nucleosomal DNA binding severely affects the structure of the underlying nucleosome, which can facilitate the subsequent binding of additional transcription factors (Ng et al., 1997). It thus appears that BZLF1 has optimized this fundamental function of AP-1 transcription factors to support EBV’s escape from repressed chromatin.

EBNA1 was also proposed to have similarities with the paradigmatic pioneer factor FoxA1 in a recent review (Niller and Minarovits, 2012), but to our knowledge, our biochemical and functional data identify BZLF1 as a *bona fide* pioneer factor of Epstein-Barr virus. Two domains in EBNA1 mimic the AT-hooks of certain cellular high mobility group proteins (Altmann et al., 2006; Hung et al., 2001) and promote the mobility of the linker histone H1 indicative of an EBNA1 intrinsic remodeling function, which is independent of cellular chromatin remodelers (Coppotelli et al., 2013).

The pioneer factor BZLF1 could share certain functions with its cousin, VP16 of Herpes simplex virus. Herpes simplex virus DNA is not (Leinbach and Summers, 1980; Muggeridge and Fraser, 1986) or only selectively associated with nucleosomes during lytic infection (Oh et al., 2015 and references therein), and partially conflicting data suggest that either histone chaperones (Oh et al., 2012), chromatin modifying co-activators (Herrera and Triezenberg, 2004), or chromatin remodelers (Neely et al., 1999) are responsible for the lack of histones on Herpes simplex DNA. Interestingly, the transactivation domain of VP16 was found to associate directly with members of the SWI/SNF remodeling complex (Neely et al., 1999) activating *in vitro* transcription from a nucleosomal template (ibid). This latter finding is reminiscent of BZLF1’s transactivation domain interacting with the chromatin remodeler INO80 (Fig. 6C), which seems to be critical for efficient lytic infection of EBV (Fig. 7).

It has also emerged from recent work (Soufi et al., 2015) that key factors in cellular reprogramming to yield induced pluripotent stem cells share critical functions with pioneer factors. Our current findings suggest that EBV has acquired this principle and puts it to use with its BZLF1 factor to reprogram viral latent chromatin within hours and to promote escape from latency.

Taken together, our experiments suggest that BZLF1 is a *bona fid*e pioneer transcription factor (Zaret and Mango, 2016), which recruits cellular machines for opening up repressed viral chromatin. It remains to be shown how BZLF1 can interact with nucleosomal DNA at high structural resolution, because BZLF1’s *in vitro* binding does not seem to follow known rules of nucleosome phasing (Fig. 4D). Future experiments should solve this conundrum.

## ACKNOWLEDGEMENT

We thank Peter Becker, Munich, for the generous usage of his fly facilities and the supply of *Drosophila melanogaster* embryos for the preparation of histone octamers. We thank Bill Sugden, Madison, for critically reading our manuscript and valuable suggestions. We are grateful to Andreas Ladurner and Karl-Peter Hopfner, Munich, for providing us with the cloned human cDNAs of *SNF2h* and *INO80*, respectively. We also thank Dagmar Pich, Munich, for precious experimental advice.

